# Hippocampal-prelimbic coupling during context-dependent extinction retrieval in rats

**DOI:** 10.1101/2025.05.03.652056

**Authors:** Flávio Afonso Gonçalves Mourão, Michael S. Totty, Tuğçe Tuna, Stephen Maren

## Abstract

After fear conditioning, repeated presentation of the conditioned stimulus (CS) alone produces a context-dependent extinction of learned fear. The hippocampus has a critical role in this process, but the mechanism by which contextual information encoded by the hippocampus leads to fear suppression is unknown. We hypothesize that contextual information encoded by the dorsal hippocampus supports the recall of extinction memory by the medial prefrontal cortex (mPFC). To test this hypothesis, we evaluated the oscillatory coherence and directional coupling of the hippocampus and mPFC during context-dependent extinction retrieval in a previously published experiment. In this experiment, male and female rats were subjected to auditory fear conditioning followed by fear extinction and extinction retrieval procedures. Previous analyses focused on oscillatory coupling during the CS; here, we performed new analyses to assess hippocampal-prefrontal coupling in the context in which extinction occurred. We found that, after extinction, re-exposing the animals to the extinction context produces a marked increase in dorsal hippocampal theta (6-8 Hz) oscillations. This increase was associated with enhanced coherence with the prelimbic (PL), but not the infralimbic (IL), division of the mPFC. Moreover, Granger causality analyses revealed that hippocampal theta oscillations preceded theta in the PL throughout the extinction retrieval session. This effect emerged during exposure to the extinction context and persisted during the presentation of the CSs and the expression of freezing behavior. Interestingly, this pattern of coherence was not observed in the IL. These results reveal that oscillatory coupling between the dorsal hippocampus and PL facilitates the context-dependent retrieval of the extinguished fear memory.

## INTRODUCTION

The ability to recall events with regard to the environmental conditions or “context” depends on associations between place, time, sensory cues, and emotional states. This process is critical for guiding behavior based on past experiences, especially in fearful and traumatic events (Maren et al., 2013).

One example of a highly context-dependent memory mechanism is extinction, a procedure often employed in behavioral therapy to treat conditions such as post-traumatic stress disorder (Vervliet et al., 2013). It relies on repeated exposure to a dangerous conditioned stimulus (CS) within a safe context, which promotes the formation of new associations that compete with the retrieval of fear memories. It is worth mentioning that this process does not erase the original fear memory; instead, it creates a new inhibitory memory that competes with the fear response when the CS is re-experienced (Bouton, 1993; Maren, 2011).

Extensive research supports the hypothesis that contextual memory encoded by the hippocampus (HPC) plays a critical role in driving fear extinction (Bouton et al., 2006; Corcoran et al., 2005; Holt & Maren, 1999; Ji & Maren, 2007; Kim & Cho, 2020; Maren & Holt, 2000). Context-dependent retrieval of extinction memories is mediated by an interconnected neural network, including the hippocampus, amygdala, and medial prefrontal cortex (mPFC) (Maren et al., 2013). Each region contributes a distinct modulatory function (Bulkin et al., 2016; Corcoran et al., 2005; Kim & Cho, 2020; Knapska et al., 2012; Plas et al., 2024; Preston & Eichenbaum, 2013; Terranova et al., 2022). For example, although both the dorsal and ventral hippocampus are involved in the modulation of contextual learning (Jin & Maren, 2015a; Maren & Holt, 2000), the dorsal hippocampus (dHPC) has a more critical role in spatial navigation (Hartley et al., 2014; O’Keefe & Nadel, 1978). In contrast, the ventral hippocampus (vHPC) plays a critical role in regulating emotional behaviors (Herman et al., 2005; Jimenez et al., 2018). Within the mPFC, the prelimbic (PL) and infralimbic (IL) regions can cooperate synergistically during the conditioned fear modulation (da Silva Vargas et al., 2021; Watanabe et al., 2021). The IL is primarily associated with facilitating fear extinction (Bloodgood et al., 2018; Jun et al., 2025; Ng et al., 2024), whereas PL is more linked to fear expression (Corcoran & Quirk, 2007; Vidal-Gonzalez et al., 2006). Though the IL and PL have often been framed as having antagonistic roles in the regulation of the inhibition and expression of fear, respectively, functional distinctions between these areas have been challenged (Giustino & Maren, 2015).

Theta oscillations are a fundamental mechanism for synchronizing neural networks during fear conditioning (Chen et al., 2021; Likhtik et al., 2014; Seidenbecher et al., 2003) and extinction (Davis et al., 2017; Lesting et al., 2013; Totty et al., 2023). In line with previous literature (Karalis et al., 2016), we have reported that after fear conditioning, the CS is associated with increased spectral power in the 3–6 Hz range in the mPFC. However, during extinction retrieval, the CS is predominantly associated with spectral power in the 6–9 Hz range in the mPFC, accompanied by significant coherence with dHPC at this oscillatory frequency range (Totty et al., 2023). However, it remains unclear whether coherence in the hippocampal-prefrontal cortex emerges before CS onset when animals use the contextual cues to recall context-appropriate memories. To address this question, we performed new analyses on a previously published data (Totty et al., 2023). In this experiment, male and female rats were subjected to auditory fear conditioning followed by fear extinction and extinction retrieval procedures. These new analyses focus on the pre-CS period when animals used contextual cues to guide memory retrieval. We found that re-exposure to the extinction context after extinction training increases 6-8 Hz theta oscillatory activity in the dHPC and increased coherence between dHPC and PL. Additionally, Granger causality analysis revealed that theta oscillations in dHPC consistently drove the PL, but not the IL, across the extinction retrieval session. These results suggest that contextual memory retrieval, encoded by the dHPC, enhances the mechanisms underlying the recall of extinction memory.

## MATERIALS AND METHODS

### Experimental procedures

The present study is based on a new analysis of previously collected local field potential (LFP) data. Portions of this dataset have been previously published with a different focus. Detailed surgical and behavioral procedure descriptions are available in the published work (Totty et al., 2023). Briefly, the animals consisted of six (3 male and 3 female) Long-Evans Blue Spruce rats that underwent an auditory fear conditioning in Context A, and context exposure, followed by extinction training and extinction retrieval task in Context B. The animals were implanted with 4×4 tungsten microwire arrays at the dHPC CA1 and 2×8 tungsten microwire arrays at the mPFC, targeting IL and PL (wires were 50-μm in diameter spaced 200-μm apart; Innovative Neurophysiology; Durham, NC). Raw signals were acquired from a monopolar design and stored using the OmniPlex system from Plexon (Dallas, TX) at 40kHz. The auditory fear conditioning task consisted of a 180-s stimulus-free baseline followed by five auditory CSs (10 s, 80 dB, 8 kHz) paired with a footshock unconditioned stimulus (US; 1.0 mA; last 2s of the CS). Over the days that followed, LFPs were recorded during a 20-min context exposure session (no CS, no US), a fear extinction session (180-s baseline; 45 CS; no US), and an extinction retrieval session (180-s baseline; 5 CS; no US), all of which occurred in the same experimental context (B) but distinct from that in which conditioning occurred (A). Each animal’s activity was measured using a load-cell platform (Med Associates Inc.) and recorded by Omniplex software from Plexon, thereby synchronizing the behavioral and electrophysiological data. Freezing behavior was defined when movement was below 5% of the maximum movement signal peak for at least one second (Figure 1).

**Figure 1:**
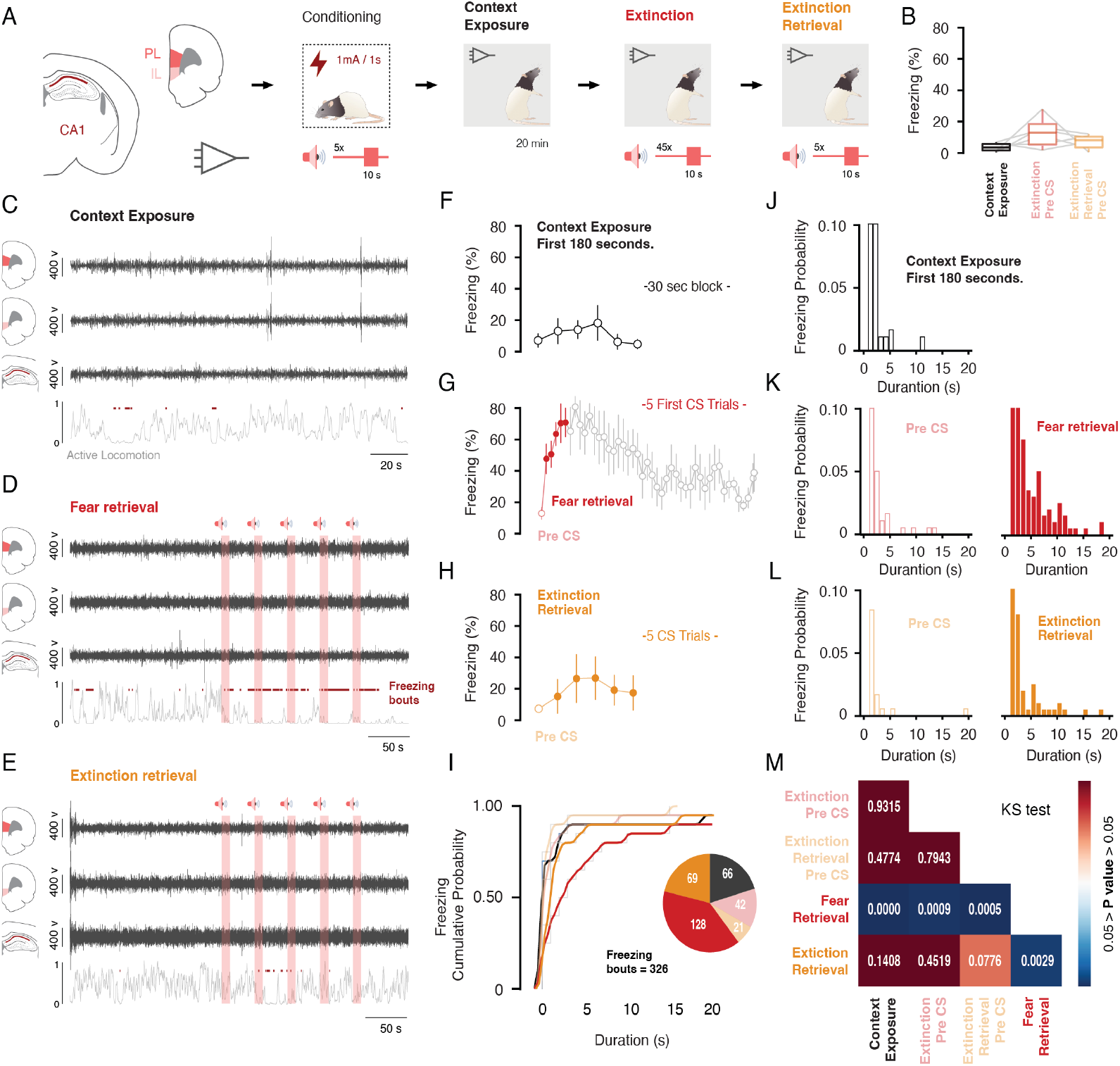
Experimental design, electrophysiological recordings, and the distribution of conditioned freezing behavior across experimental sessions. **A**. Neural substrates recorded and the behavioral timeline of the fear conditioning protocol, as outlined by Totty et al. (2023). **B**. Percentage of freezing measured throughout the pre-CS period at each session. **C, D, E**. Representative filtered signals (2-12 Hz) from mPFC-PL, mPFC-IL, and dHPC during each experimental session. Red-light shaded areas indicate the presentation of the conditioned stimulus (CS), while red dots mark freezing episodes. The light gray line below the LFP time course represents movement activity, measured using the load-cell system (Med-Associates Inc.). **F, G, H**. Percentage of freezing measured throughout each experimental session. The context exposure sample was divided into 30-second time windows (totaling 180 seconds). During extinction, freezing was measured in the pre-CS period followed by 45 CS trials, with the fear retrieval period highlighted in red (first five trials). Extinction retrieval included the pre-CS period and five CS trials. **J, K, L**. Probability distribution of freezing bout durations for context exposure, extinction pre-CS, fear retrieval, extinction retrieval pre-CS, and extinction retrieval CS presentations. **I**. Cumulative probability function of freezing bouts across each experimental session. The pie chart represents the total number of freezing bouts per experimental session. **M**. A two-sample Kolmogorov–Smirnov test (KS test) was used to assess whether freezing bouts came from the same distribution, comparing each experimental phase/session.

### Data analysis

All the analyses presented here focus on the ∼180-s stimulus-free period preceding the presentation of the CS (i.e., pre-CS period). During this period, animals rely on recognition memory to determine if they are in a novel or familiar context, and this, in turn, directs context-dependent memory retrieval processes. We hypothesized that hippocampal-prefrontal coupling during this period is critical for the suppression of conditioned freezing to an extinguished CS in the extinction context. To assess this, we examined hippocampal-prefrontal coherence during this pre-CS period in three behavioral sessions: 1) context exposure (i.e., exposure to a novel context that would ultimately host extinction), 2) extinction, and 3) extinction retrieval. Specifically, spectral power analyses and coherence between the dHPC and mPFC (PL and IL) theta oscillations (2-12 Hz) were examined during the 180-second pre-CS period of each of these sessions. Additionally, Granger causality analyses between the dHPC and mPFC (PL and IL) were performed during the presentation of the CSs and throughout the inter-trial intervals (ITIs) (one animal was excluded due to large movement artifacts during the extinction retrieval phase, which compromised the autoregressive model). All analyses were performed using built-in and custom codes in MATLAB (https://github.com/marenlab). LFPs from one single channel were obtained by removing the linear tendencies for a constant mean and down-sampling to 1 kHz. Periods with motion artifacts and/or line noise were removed from all analyses.

#### Power spectral density and time-frequency power

Power spectra and illustrative time-frequency power spectrograms were calculated using the *pwelch*.*m* and *spectrogram*.*m* functions, respectively (Signal Processing Toolbox). Power spectra were set as 2,048-point, and time-frequency power spectrograms were set as 30,000-point, Hamming window. For both functions, *nfft* = 2^15^ and 90% overlapping were used. Power spectra density and spectrograms were normalized to the total power between 2-12 Hz. The average power between 6-8 Hz was then calculated to obtain a power estimate.

#### Spectral coherence

The spectral coherence between the correlated brain regions was calculated using the *mscohere*.*m* function with a 2,048-point, Hamming window, 90% overlapping and *nfft* = 2^15^. The average spectral coherence between 6-8 Hz was then calculated. Illustrative coherograms were generated using the *MTCoherogram*.*m* function, which employs multi-taper estimation based on Chronux software (http://chronux.org). This function was adapted from the Freely Moving Animal (FMA) Toolbox developed by Michaël Zugaro (https://fmatoolbox.sourceforge.net/), set as 18,000-point, 95% overlapping and relative resolution and order of the tapers as 3 and 5, respectively.

#### Phase lock value

The data were initially filtered within the frequency range of 6-8 Hz using the *eegfilt*.*m* (EEG lab; https://sccn.ucsd.edu/eeglab/). The phase coefficients were extracted using the *hilbert*.*m* function (Signal Processing Toolbox™). The Δ phase was calculated in 250 ms average time windows as the difference between the imaginary components of the correlated brain regions.

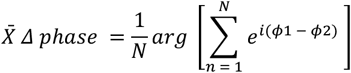

Where 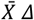 *phase* is the argument of the sum of the phase vectors; N is the number of time-axis samples of each signal; ϕ1 and ϕ2 are the phase values for the brain regions. Phase lock value (PLV) is a measure of phase coherence represented by a dimensionless real value ranging from 0 to 1. It was quantified as the magnitude of the mean vector derived from the differences in phase angles. A value of 0 indicates a uniform phase distribution, while a value of 1 signifies perfect phase alignment (Lachaux et al., 1999).

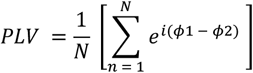

#### Granger causality

The Wiener-Granger causality method (Barnett & Seth, 2014) was employed to evaluate whether the past activity of one time series can predict the future activity of another, providing insight into the causal relationships underlying neural interactions. Initially, the LFPs were *z*-scored and downsampled to 250 Hz. The model order was calculated using the Levinson-Wiggins-Robinson (LWR) algorithm. For the pre-CS, CS, and ITI periods, the regression model was associated with the Bayesian Information Criterion (BIC). The model order was set to ∼25, which provided a good fit to the data. This choice is also consistent with previous findings, which reported that longer time windows enhance the reliability of model estimation (de Almeida-Filho et al., 2021). Pairwise conditional spectral Granger causality was computed using the *var_to_spwcgc*.*m* function, and the value across the desired frequency range (6-8 Hz) was calculated by averaging the Granger spectrum with the *smvgc_to_mvgc*.*m* function. Finally, a statistical control analysis was performed for each pairwise condition by surrogating the time series 1000 times using the *permtest_tsdata_to_pwcgc*.*m* function. Illustrative Granger coherograms were generated using sliding time windows across the task (20 s with 90% overlap). The Granger spectrum for each time epoch was calculated with the *var_to_spwcgc*.*m* function, as described above.

### Statistics

All data were expressed as means ± standard error of the mean (SEM). The approximation to the normal distribution was confirmed by the Kolmogorov– Smirnov test (*p* > 0.05). Statistical comparisons were made using repeated-measures analysis of variance (ANOVA), followed by Tukey’s multiple comparisons test. Values of *p* < 0.05 were considered statistically significant. To evaluate whether the empirical distributions of freezing bouts across experimental phases came from the same distribution, the Kolmogorov-Smirnov test for two samples was applied (*p* < 0.05). Data were analyzed using GraphPad Prism 9.0 software and MATLAB 2022a (The Mathworks, Natick, United States).

## RESULTS

### The probability of conditioned responses during extinction retrieval resembled baseline levels from pre-CS and context exposure sessions, indicating reduced CS-driven freezing behavior

Behavioral data from these experiments were previously published (Totty et al., 2023). Here, we present additional analyses aimed at further characterizing the statistical properties of freezing bouts across experimental sessions. First, we aimed to rule out the possibility that the differences in LFPs across experimental sessions were due to differential freezing levels during these experimental sessions. Specifically, we tested the hypothesis that freezing behavior during the pre-CS period was comparably low across sessions, suggesting that animals were in a similar behavioral state prior to cue onset. Second, we aimed to demonstrate that the conditioned freezing during CS presentations in the extinction retrieval session is similar to the freezing behavior during the pre-CS period, suggestive of successful extinction retrieval (Figure 1F-M). To this end, we assessed the probability distribution function of conditioned freezing responses as a function of their duration (Figure 1J-L) following a cumulative distribution analysis (Figure 1I-M). This approach revealed that freezing levels were similar during context exposure, extinction, and extinction retrieval pre-CS periods (Figure 1B). In addition, during the extinction retrieval session, the distribution curve of the conditioned response closely resembled the probability observed over the pre-CS and context exposure periods (Figure 1I & J-M). Furthermore, as expected, during the fear retrieval phase (first five trials of extinction), the cumulative distribution curve was significantly shifted to the right, due to a higher density of longer lasting freezing bouts (Kolmogorov–Smirnov test for two samples, *p* < 0.05; Figure 1M). These results confirm that the pre-CS freezing levels during context exposure, extinction, and extinction retrieval and CS freezing levels during extinction retrieval were comparable. Therefore, differences in LFP signal cannot be explained by differences in conditioned behavior as measured by locomotor activity.

### Context reinstatement enhances theta activity in the dHPC and its synchronization with the mPFC-PL

Hippocampal theta oscillations are essential for contextual memory, insofar as they synchronize neural activity across brain regions involved in memory encoding and retrieval (Buzsáki and Moser, 2013; Staudigl and Hanslmayr, 2013; Belchior et al., 2014). Theta oscillations are thought to support the integration of information with its environmental context during encoding (Wirt & Hyman, 2019). Enhanced theta power during encoding has been associated with improved subsequent retrieval, particularly when contextual cues are reinstated (Shrager et al., 2008). These findings suggest that theta rhythms play a critical role in forming and accessing episodic memories by associating them with the context in which they were initially acquired (Herweg et al., 2020; Staudigl & Hanslmayr, 2013). Accordingly, to test the hypothesis that recognition of the familiar context during the extinction retrieval session would be accompanied by an increase in theta activity, we performed spectral analyses on neural recordings throughout the pre-CS period. As shown in Figure 2 (J, K), during the extinction retrieval pre-CS period, the dHPC displayed a significant increase in normalized 6-8 Hz power (*F*_2,10_ = 12.12, *p* < 0.01) compared to the extinction pre-CS (post hoc: *p* < 0.05) and the context exposure period (*p* < 0.0001). No significant differences were found in normalized 6-8 Hz power when assessed in the mPFC-PL (Figure 2D, E) and mPFC-IL (Figure 2G, H).

**Figure 2:**
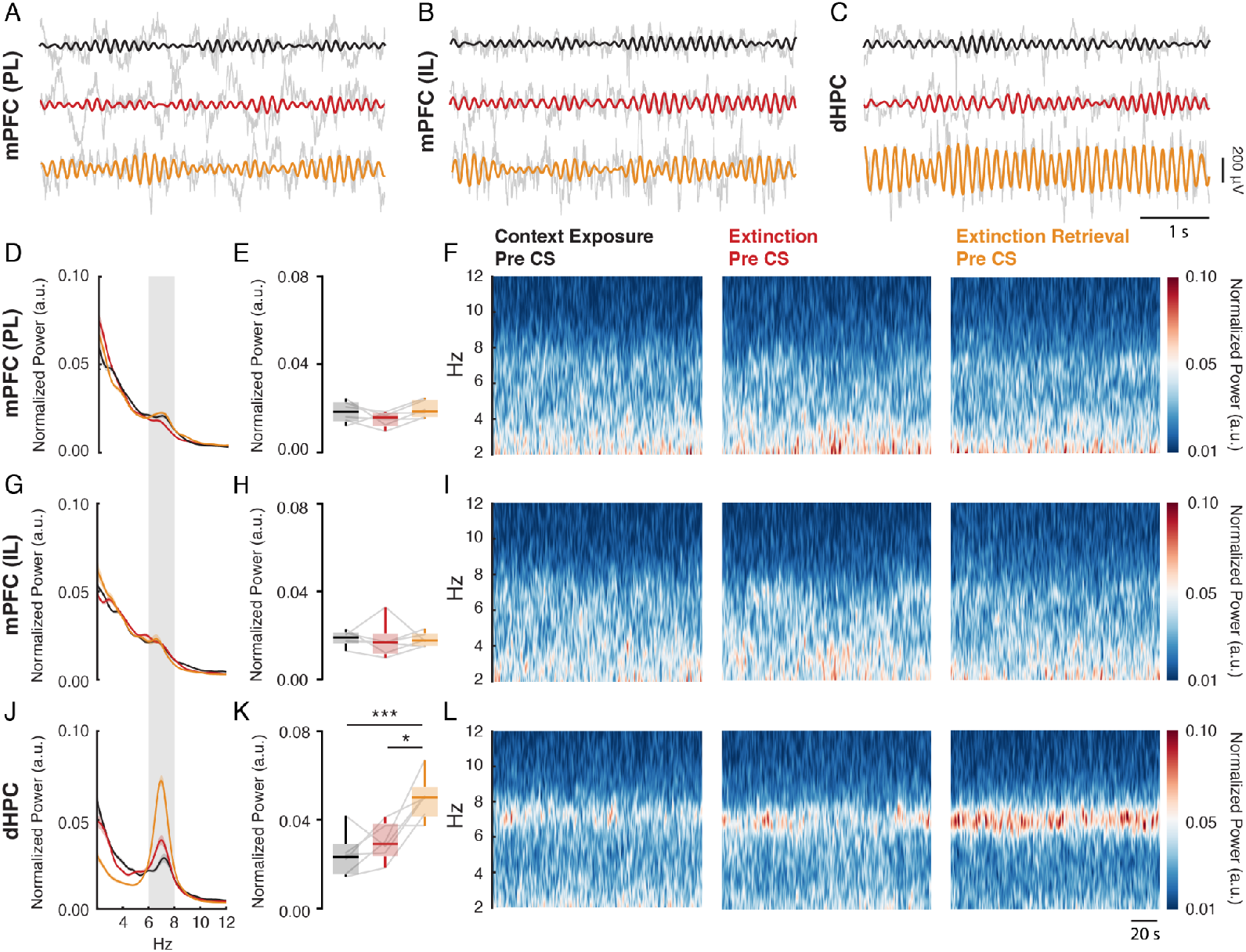
Dorsal hippocampus theta power increases when animals are re-exposed to the extinction context during the extinction retrieval session. **A, B, C**. Representative time courses from mPFC-PL, mPFC-IL, and dHPC during the pre-CS period. The raw signal is shown in gray, while the filtered signals at 6-8 Hz for context exposure, extinction, and extinction retrieval pre-CS periods are shown in black, red, and orange, respectively. **D, G, J**. Normalized spectral power density between 2-12 Hz from mPFC-PL, mPFC-IL, and dHPC. **E, H, K**. During the extinction retrieval pre-CS, the dHPC exhibited increased normalized 6-8 Hz power compared to the context exposure (*p* < 0.0001) and the extinction pre-CS (*p* < 0.05). **F, I, L**. Time-frequency spectrograms between 2-12 Hz from mPFC-PL, mPFC-IL, and dHPC during the pre-CS period.

Although evidence suggests that repeated exposure to familiar contexts can reduce exploratory behavior, consistent with a typical habituation response to a known environment (Leussisa & Bolivar, 2006), other findings indicate that rodents in unfamiliar environments tend to move at slower speeds, reflecting a more cautious approach to novel surroundings. As familiarity increases, their locomotor speed typically rises, suggesting confidence in navigation (Arkley et al., 2014). Given that changes in locomotor activity have a substantial influence on hippocampal theta activity (Belchior et al., 2014; Bender et al., 2015; Kropff et al., 2021; McFarland et al., 1975), we performed a complementary analysis focusing on the animals’ movement dynamics and the correlations between the relative amplitude envelope (6-8 Hz) and the active locomotion. We found that the probability distribution function of activity bouts was similar (Supplementary Figure 1), and theta activity was significantly correlated with locomotor measurement (Supplementary Figure 2) across all pre-CS periods. This suggests that motor activity during contextual presentations is unlikely to be a primary factor contributing to the observed results.

Furthermore, as shown in Figure 3 (A-C), the pre-CS period during the extinction retrieval test was accompanied by increased spectral coherence between dHPC and mPFC-PL (*F*_2,10_ = 8.48, *p* < 0.01) compared to the pre-CS period during the extinction session (*p* < 0.05) and the context exposure session (*p* < 0.01). Because spectral coherence also includes power information, the results of this analysis are likely to be influenced by significant increases in power. If connectivity increases without a corresponding change in power, as observed in the mPFC, spectral coherence may yield biased results (Cohen, 2014). Based on this, we also conducted an analysis of the relative phase coherence between the dHPC and both the PL and IL, aiming to understand how these substrates interact at a dynamic and temporal level (Lachaux et al., 1999; Womelsdorf et al., 2007). Consistent with previous findings, the enhanced spectral synchronization was further corroborated by a notable alignment of relative phase within the 6-8 Hz range between the dHPC and mPFC-PL (Figure 3G-I). In particular, extinction retrieval pre-CS was associated with significantly higher phase coherence (PLV) (*F*_2,10_ = 7.72, *p* < 0.01) compared to both the extinction pre-CS (*p* < 0.05) and the context exposure period (*p* < 0.01). Regarding the mPFC-IL, although spectral coherence was not significant (Figure 3D-F), phase coherence (Figure 3J-L) during extinction retrieval pre-CS was significantly higher compared to the context exposure period (*F*_2,10_ = 5.63, *p* < 0.05).

**Figure 3:**
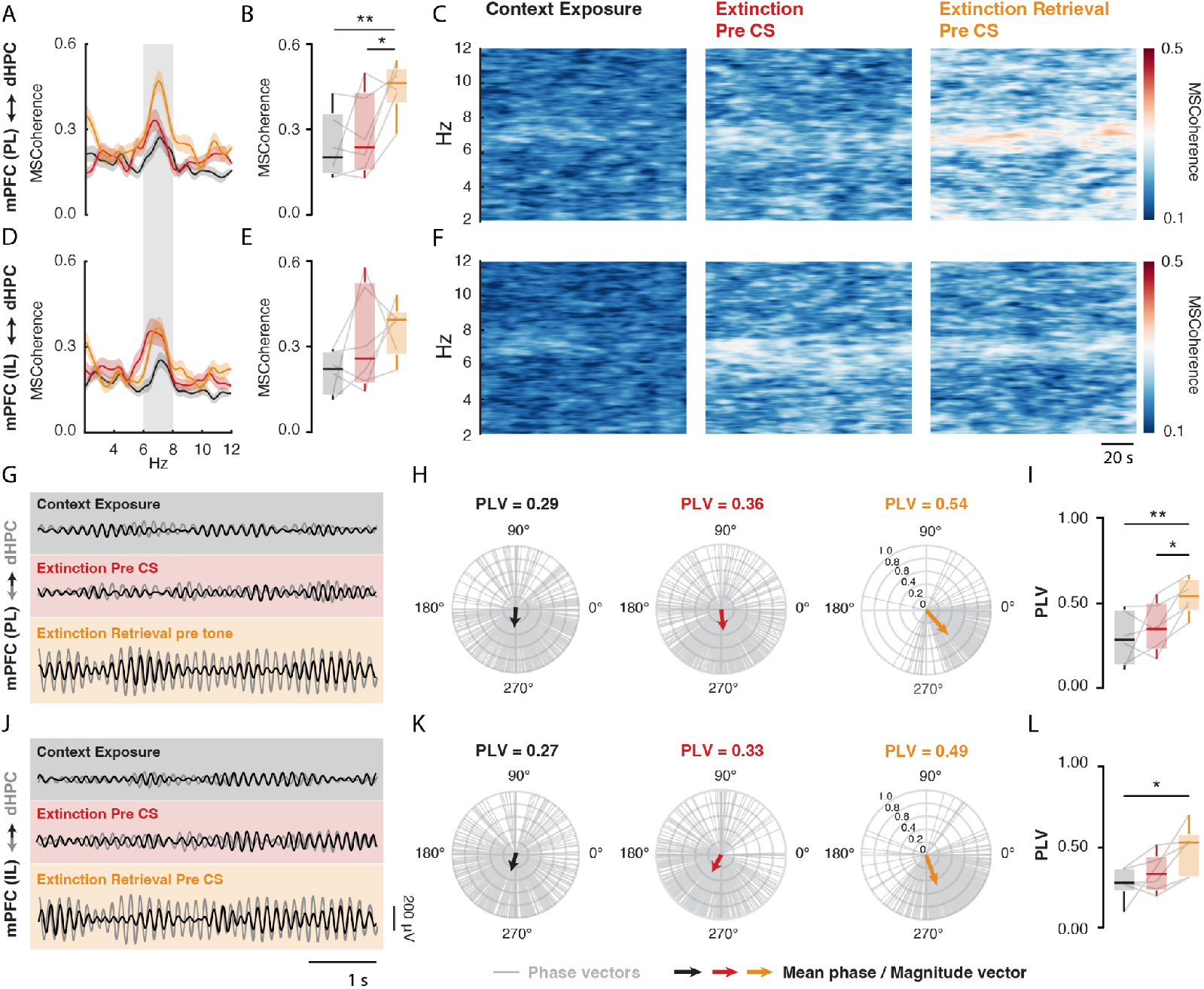
mPFC and dHPC theta oscillations increase synchrony when animals are re-exposed to the extinction context during the extinction retrieval session. **A, D**. Magnitude-squared coherence (2-12 Hz) between dHPC and mPFC-PL (**A-B**), as well as dHPC and mPFC-IL (**D-E**), during the pre-CS period. **B**. Over the extinction retrieval pre-CS period, the dHPC and mPFC-PL exhibited increased coherence at 6-8 Hz compared to the first context exposure (*p* < 0.001) and extinction pre-CS (*p* < 0.05). Time-frequency coherograms between 2-12 Hz from dHPC and mPFC-PL (**C**), as well as dHPC and mPFC-IL (**F**) during the pre-CS period. **G, J**. Representative overlapping time courses of filtered signals (6-8 Hz) for context exposure, extinction, and extinction retrieval pre-CS periods. **H, K**. Delta phase vectors computed every 250 ms at 6-8 Hz during the pre-CS period (gray lines), with the corresponding estimated mean phase values represented by arrows. The numbers above the polar plots indicate the phase-locking value (PLV) for each period. **I, L**. During the extinction retrieval pre-CS period, the dHPC and mPFC-PL exhibited increased values at 6-8 Hz compared to the context exposure (*p* < 0.0001) and extinction pre-CS (*p* < 0.05), while the dHPC and mPFC-IL exhibited increased values compared to the first context exposure (*p* < 0.05).

### dHPC theta oscillations drive the mPFC-PL activity during the extinction retrieval session

Evidence suggests that the interaction between the HPC and mPFC is bidirectional, reflecting distinct yet cooperative mechanisms for memory representation (Jin & Maren, 2015b; Place et al., 2016; Preston & Eichenbaum, 2013). Overall, functional connectivity analyses revealed a flow of contextual information from the HPC to the mPFC during contextual memory retrieval during the extinction retrieval session (Place et al., 2016). To test whether dHPC theta oscillations predict activity in the mPFC during extinction retrieval, we performed Granger causality analysis (Barnett & Seth, 2014). This approach allows us to determine whether the observed synchrony reflects a hierarchical relationship, suggesting an active role of the dHPC in influencing the mPFC (Seth et al., 2015).

Granger causality analysis revealed that, during the pre-CS period of the extinction retrieval session, dHPC theta oscillations predict the mPFC-PL oscillations in the 6-8 Hz range (*F*_2,8_ = 5.47, *p* < 0.05), with significant differences compared to both the pre-CS period during the extinction session (*p* < 0.05) and the context exposure session (*p* < 0.01) (Figure 4A-B). Furthermore, significant values were found in the direction from the dHPC to mPFC-PL when compared to the opposite direction, from the mPFC to dHPC (*p* < 0.01). In contrast, no significant differences were observed in the dynamics between the dHPC and mPFC-IL across the experimental sessions (Figure 4C-D).

**Figure 4:**
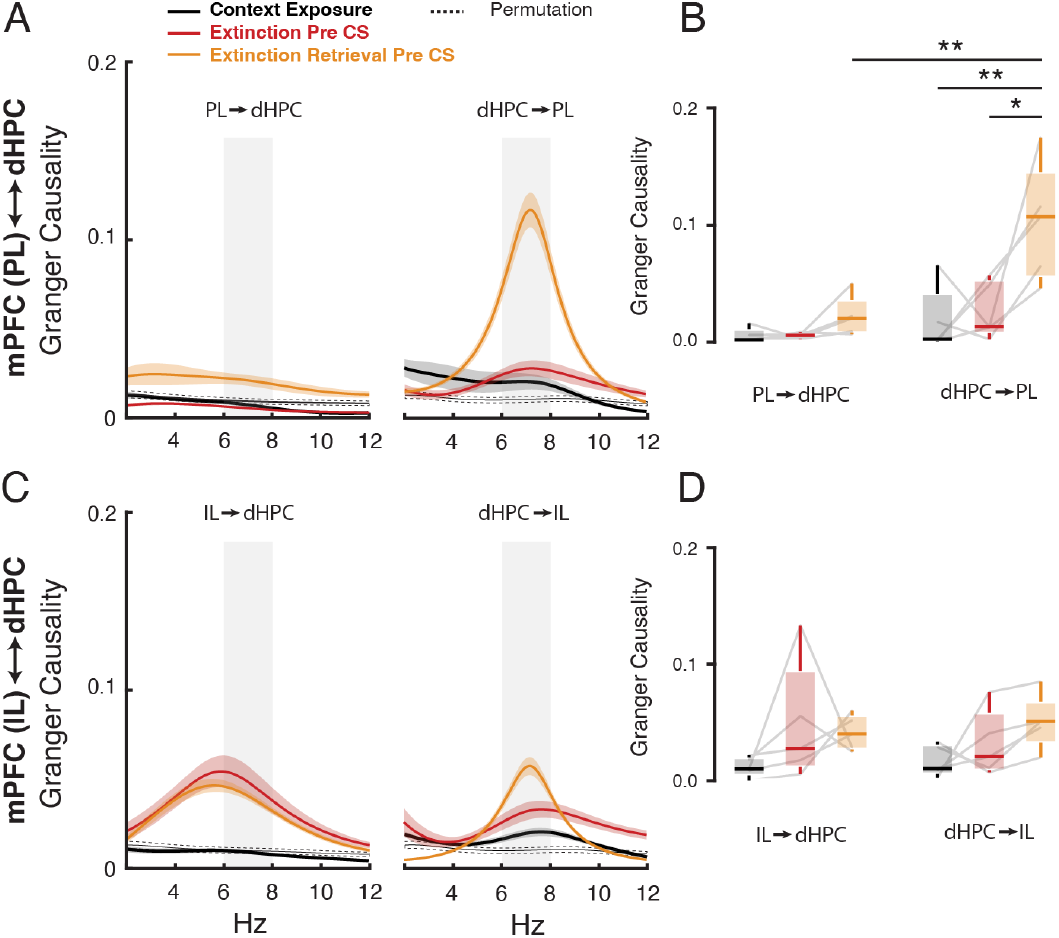
dHPC theta oscillations drive the mPFC-PL activity when animals are re-exposed to the context during the extinction retrieval session. **A**. dHPC ←→ mPFC-PL spectral Granger causality between 2 - 12 Hz during the pre-CS period. **B**. dHPC → mPFC-PL and mPFC-PL → dHPC at 6-8 Hz. The dHPC drive activity in mPFC-PL at 6-8 Hz compared to the context exposure (*p* < 0.01) and extinction pre-CS (*p* < 0.05). Furthermore, it exhibits a stronger response compared to the opposite direction from mPFC-PL → dHPC (*p* < 0.01). **C**. dHPC ←→ mPFC-IL spectral Granger causality between 2 - 12 Hz. **D**. There are no significant differences between dHPC ←→ mPFC-IL across the experimental phases.

Notably, as shown in Figure 5 (A-D), the predictive influence of dHPC theta oscillations on the mPFC-PL during the extinction retrieval session remained elevated and stable after the pre-CS phase, with a significant main effect against fear retrieval during both the CS presentation and the ITI (*F*_1,4_ = 17.24, *p* < 0.05). Furthermore, it showed a significant effect when compared to the opposite direction, from the mPFC-PL to dHPC (*F*_1,4_ = 8.49, *p* < 0.05).

**Figure 5:**
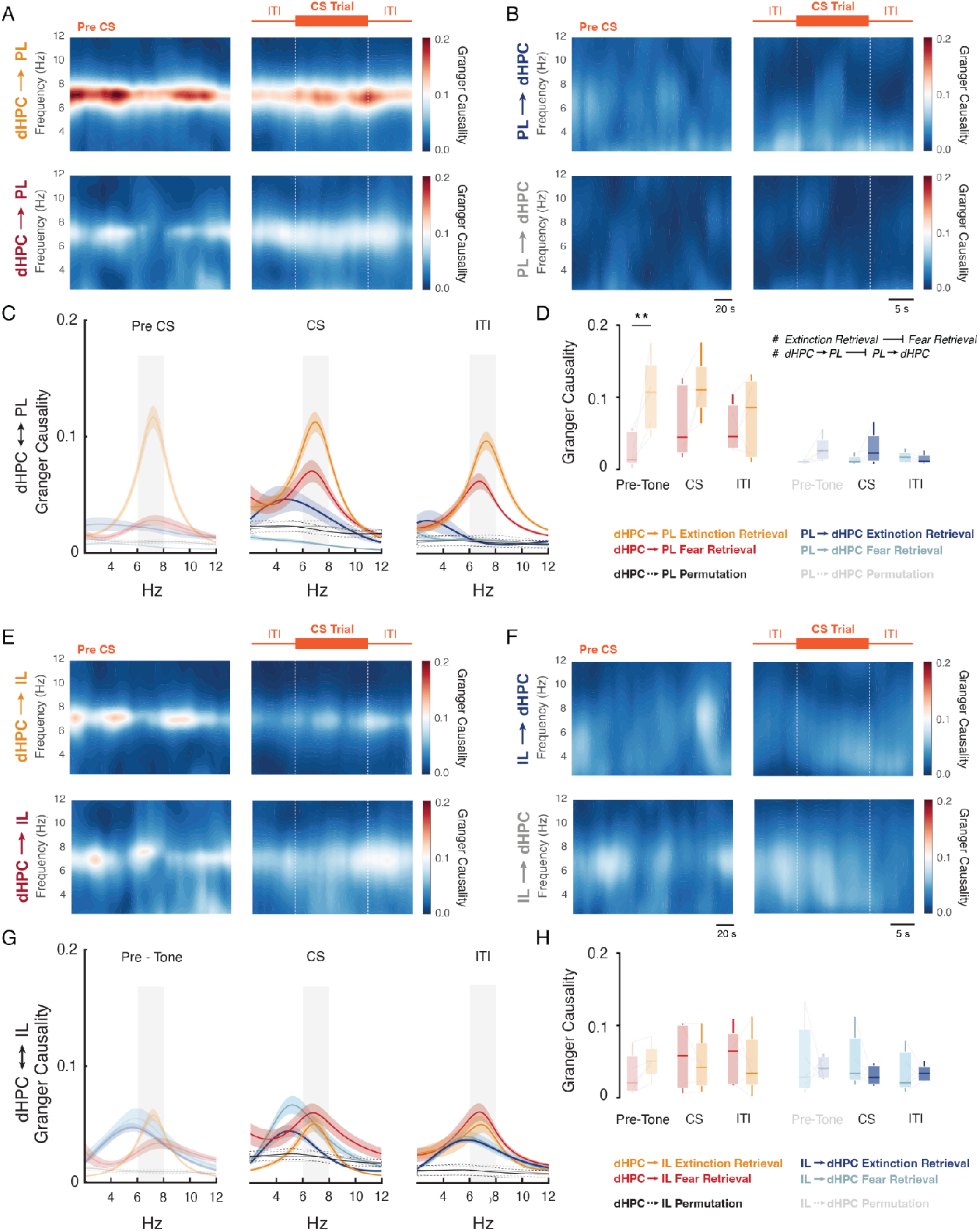
dHPC theta oscillations consistently drive the mPFC-PL activity across the extinction retrieval session. **A, B**. Time-frequency Granger causality between 2 - 12 Hz from dHPC → mPFC-PL and mPFC-PL → dHPC, respectively, for the extinction (fear retrieval) and the extinction retrieval session. **C**. dHPC ←→ mPFC-PL spectral Granger causality between 2 - 12 Hz for the extinction and the extinction retrieval session. **D**. The dHPC drives activity in the mPFC-PL at 6–8 Hz, as indicated by the analysis of variance, which revealed a significant difference between extinction and fear retrieval trials (#*p* < 0.05). Additionally, there was a significant main effect in the direction of causality, with dHPC → mPFC-PL differing from mPFC-PL → dHPC (#*p* < 0.05). **E, F**. Time-frequency Granger causality between 2 - 12 Hz from dHPC → mPFC-IL and mPFC-IL → dHPC, respectively, for the extinction and the extinction retrieval sessions. **G**. dHPC ←→ mPFC-IL spectral Granger causality between 2 - 12 Hz for the extinction and the extinction retrieval session. **H**. There are no significant differences between dHPC ←→ mPFC-IL across the experimental phases.

Ultimately, in line with previous findings, no significant differences were observed in the dynamics between the dHPC and mPFC-IL in either direction (Figure 5, E-H). These data reveal that context-dependent memory retrieval during extinction retrieval is driven by coherent activity in the dHPC-PL network, and that the oscillatory correlates of contextual recall in the dHPC influence cue-driven memory retrieval processes by the mPFC.

## DISCUSSION

After fear conditioning, presentations of the CS alone during extinction result in a context-dependent memory: fear to the CS is reduced in the extinction context but high everywhere else (Bouton, 1988; Harris et al., 2000; Bouton et al., 2021). Our new findings demonstrate that context-dependent memory retrieval is indexed by oscillatory correlates in the dHPC-mPFC network. Specifically, the dHPC maintains significant and sustained theta activity when animals are re-exposed to the familiar context after the extinction session. Furthermore, we found greater network coherence between the dHPC and mPFC-PL, which may represent strengthened functional connectivity between these regions. An increase in theta activity reflects stronger or more synchronized neural oscillations, which can reflect enhanced neural coordination and communication, often linked to encoding and retrieval of memories (Buzsáki, 2002). Furthermore, the new analyses we report here reveal that contextual memory encoded by the dHPC plays an essential role in modulating extinction memory expression, promoting discrimination between familiar contexts while allowing appropriate generalization when contexts share relevant features.

As strongly suggested by the literature, extinction memories are highly sensitive to context, as context provides cues that link salient sensory inputs and memory traces, helping to determine when a relevant behavior should be triggered (Maren et al., 2013). In this sense, our findings show that contextual memory retrieval, processed by the dHPC, directly influences mPFC dynamics. This influence likely reflects parallel processing that signals safety prior to CS presentation, ultimately impacting the success as well as the updating and maintenance of extinction memory recall (Corcoran et al., 2005; Rossato et al., 2010). It is important to highlight that this involvement can also extend to the retrieval of fear memory itself (Holt & Maren, 1999; Ye et al., 2017).

Previous evidence has led to the proposal of a bidirectional interaction model between HPC and mPFC during memory retrieval. In this framework, the HPC conveys contextual information to the mPFC, which in turn facilitates and guides successful retrieval processes (Jin & Maren, 2015b; Place et al., 2016; Preston & Eichenbaum, 2013). In line with this, we demonstrate that, upon reexposure to the extinction context, dHPC theta activity leads the mPFC-PL theta activity, suggesting a directional flow of information from the dHPC to the mPFC-PL. This might convey the safety of the extinction context to the mPFC, which might be necessary for extinction retrieval. Moreover, the CS presentations after the tone-free baseline may then trigger a reversal in information flow, where the mPFC takes the lead in driving hippocampal activity (Lesting et al., 2013; van Bree et al., 2025). This may suggest distinct yet cooperative mechanisms between the HPC and mPFC in memory representation and retrieval (Place et al., 2016). However, Granger causality analysis revealed that the dHPC theta oscillations consistently predicted those in mPFC-PL throughout the entire extinction retrieval session. This may indicate a more prolonged contextual information transfer from the HPC to the mPFC during memory retrieval than initially proposed. It is important to note that, although the HPC is the primary generator of theta oscillations in the rodent and human brain (Colgin, 2016; Shin et al., 2005; López-Madrona et al., 2020; Colgin, 2016; Shin et al., 2005) — and theta activity observed in the mPFC is largely driven by hippocampal input (Eichenbaum, 2017) — what stands out in our findings is that this dominance becomes particularly evident during the extinction retrieval session. Notably, this effect does not appear to be solely driven by contextual familiarity, as the extinction session also occurred in a familiar environment, immediately following the context exposure session. We hypothesize that the learned experience during repeated exposures to the CS, which led to the formation of the extinction memory, was strongly supported by the salience of the contextual experience itself, thereby facilitating successful retrieval of the extinction memory (Bulkin et al., 2016; Corcoran et al., 2005; Harris et al., 2000; Ji & Maren, 2007).

Previous work from our group demonstrated that the mPFC-PL and mPFC-IL networks show increased 3-6 Hz power at the onset of fear retrieval during CS presentation. During extinction retrieval, on the other hand, increased 6–9 Hz power is observed in the mPFC (Totty et al., 2023). This shift in oscillatory dynamics during extinction retrieval demonstrates that theta oscillations can reflect distinct neural states driven by pathway-specific interactions (Goyal et al., 2020; López-Madrona et al., 2020; Zheng et al., 2025). Here, we additionally showed an increase in 6-8 Hz power in the dHPC upon exposure to the familiar extinction context. These hippocampal theta oscillations, mainly synchronized with those of the mPFC-PL but also with the IL — as evidenced by a well-defined phase relationship — may be critical for extinction retrieval. Consistent with this, the thalamic nucleus reuniens, a critical hub for communication between the HPC and the mPFC, actively modulates mPFC-HPC oscillatory activity during extinction retrieval (Ferraris et al., 2021; Hauer et al., 2019; Plas et al., 2024; Silva et al., 2021; Totty et al., 2023). Moreover, inhibition of the nucleus reuniens during extinction retrieval decreases the HPC-mPFC oscillatory coherence at 6-9 Hz, leading to poor extinction retrieval (Ramanathan et al., 2018; Totty et al., 2023).

Within a broader functional framework, the scientific community has increasingly explored causal relationships between particular patterns of neural oscillations and biological functions (van Bree et al., 2025; Zheng et al., 2025). While this is a promising direction, we believe that caution is warranted when making definitive claims. Although oscillations are a hallmark of neural circuit activity, the same oscillatory patterns should not be attributed to a single function or cognitive process. For example, mPFC activity is well-known to exhibit strong oscillations in the 3–6 Hz range during fear retrieval, which are synchronized with the amygdala (Karalis et al., 2016). However, evidence indicates that this slow-frequency range is not exclusively related to fear retrieval (Tavares and Tort, 2022). Additionally, the directionality previously observed between the mPFC-IL and dHPC during freezing behavior within this frequency range (Lesting et al., 2013) also appears to be strongly influenced by respiratory-related oscillations (Bagur et al., 2021).

It is important to highlight that anatomical specializations and interconnected neural pathways may have distinct modulatory functions (Fanselow & Dong, 2010; Sierra-Mercado et al., 2011; Vidal-Gonzalez et al., 2006). For example, vHPC projects to inhibitory interneurons in the mPFC-IL, resulting in feed-forward inhibition. Activation of this pathway is associated with the renewal of previously extinguished fear, playing a crucial role in the relapse of fear responses (Marek et al., 2018). On the other hand, the dHPC, specifically the CA1/subiculum region, exerts a direct excitatory influence on interneurons in the mPFC-PL and medial-orbital areas. A significant portion of this excitatory pathway is likely monosynaptic, with its direct output being dynamically targeted through this inhibitory process. Consequently, this could enhance the synchronization of a specific subset of mPFC-PL and medial-orbital neurons with dHPC activity (Tierney et al., 2004). Also, the HPC modulation of mPFC activity may influence amygdala output, thereby contributing to the regulation of fear expression in response to the CS (Maren, 2011; Plas et al., 2024; Sierra-Mercado et al., 2011).

Finally, although the new analyses suggest a mechanism by which dHPC modulates mainly mPFC-PL dynamics enhancing the retrieval of extinction memories when a familiar context is re-encountered, additional experimental controls are needed to support this hypothesis — for example, groups exposed only to the context across days without the US, or groups receiving unpaired presentations between CS-US. Nonetheless, it is important to note that previous evidence indicates that re-exposure to a context following salient events is associated with increased hippocampal activity, which in turn predicts the strength of memory retrieval (Shrager et al., 2008).

In conclusion, the present study suggests that the dynamic interaction between the dHPC and the mPFC-PL may represent a potential mechanism underlying the retrieval of extinction memory across familiar contexts.

## AUTHOR CONTRIBUTIONS

FM analyzed the behavioral and electrophysiological data. MT performed the experiments. FM, TT, MT and SM wrote the manuscript.

## DATA AVAILABILITY

The data from these experiments are available from the corresponding author upon request

## CODE AVAILABILITY

Custom codes used for data analyses are available at https://github.com/marenlab.

## ACKNOWLEDGEMENTS

We thank Dr. Eduardo Mazoni Andrade Marçal Mendes from Federal University of Minas Gerais, Brazil, and Vinícius Rezende Carvalho from University of OSLO, Norway, for insightful discussions on the code and valuable guidance in interpreting the signal processing analyses.

## SUPPLEMENTARY MATERIAL

**Supplementary Figure 1:**
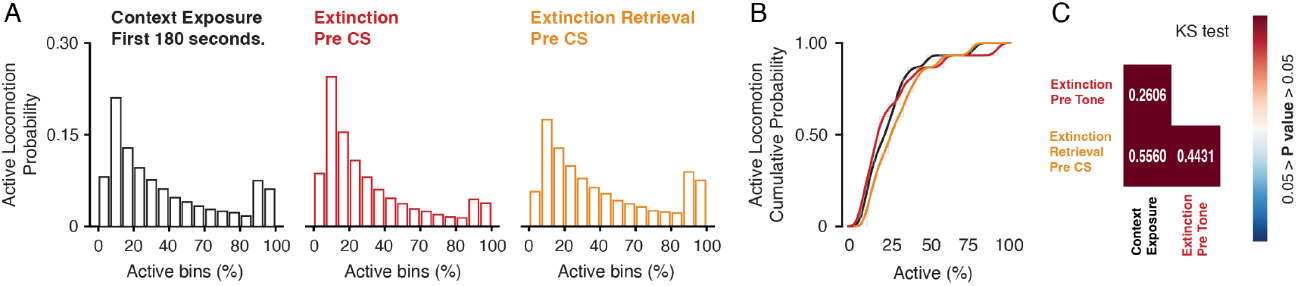
Probability distribution function of active locomotion bouts across experimental sessions. **A**. Probability distribution of active locomotion bouts for context exposure, extinction pre-CS, and extinction retrieval pre-CS **B**. Cumulative probability function of active locomotion bouts across each experimental session. **C**. A two-sample Kolmogorov–Smirnov test (KS test) was used to assess whether active locomotion bouts came from the same distribution, comparing each experimental session.

**Supplementary Figure 2:**
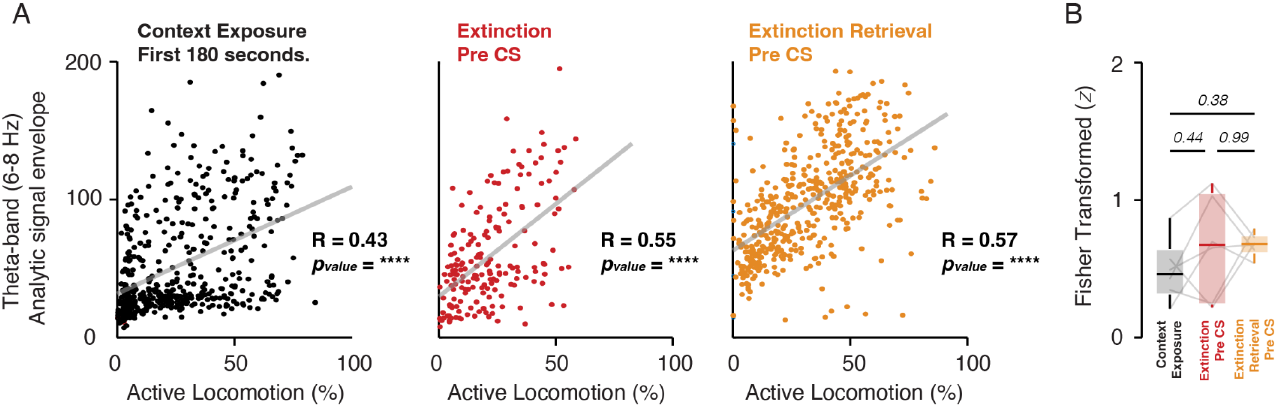
Pearson correlations between the relative magnitude envelope (6-8 Hz) and the active locomotion over the experimental sessions. **A**. Over the pre-CS period, data from all animals were extracted in sliding windows and averaged every 2 seconds. Theta-band magnitude was computed by applying the Hilbert transform to the bandpass-filtered signal (6–8 Hz), resulting in a complex analytic signal whose amplitude envelope corresponds to its magnitude. **B**. Pearson correlations were computed individually for each experimental animal. The resulting coefficients were transformed using the Fisher z-transformation, which stabilizes the variance and renders the distribution approximately normal. The transformed values were then compared using analysis of variance (ANOVA), followed by pairwise comparisons with Tukey’s post hoc test. *P-values* are indicated in the figure.

